# The TcdE holin drives toxin secretion and virulence in *Clostridioides difficile*

**DOI:** 10.1101/2023.09.16.558055

**Authors:** NV DiBenedetto, M Oberkampf, L Cersosimo, V Yeliseyev, L Bry, J Peltier, B Dupuy

**Affiliations:** Massachusetts Host-Microbiome Center, Dept. Pathology, Brigham and Women’s Hospital, Harvard Medical School, Boston, MA, USA; Institut Pasteur, Université Paris-Cité, UMR-CNRS 6047, Laboratoire Pathogenèse des Bactéries Anaérobies, F-75015 Paris, France; Institute for Integrative Biology of the Cell (I2BC), CEA, CNRS, Université Paris-Saclay, Gif-sur-Yvette, France

## Abstract

*Clostridioides difficile* is the leading cause of healthcare associated infections. The Pathogenicity Locus (PaLoc) toxins TcdA and TcdB promote host disease. These toxins lack canonical N-terminal signal sequences for translocation across the bacterial membrane, suggesting alternate mechanisms of release, which have included targeted secretion and passive release from cell lysis. While the holin TcdE has been implicated in TcdA and TcdB release, its role *in vivo* remains unknown. Here, we show profound reductions in toxin secretion in *tcdE* mutants in the highly virulent strains UK1 (epidemic ribotype 027, Clade 3) and VPI10463 (ribotype 087, Clade 1). Notably, *tcdE* deletion in either strain rescued highly susceptible gnotobiotic mice from lethal infection by reducing acute extracellular toxin to undetectable levels, limiting mucosal damage, and enabling long-term survival, in spite of continued toxin gene expression in *tcdE* mutants. Our findings confirm TcdE’s critical functions *in vivo* for toxin secretion and *C. difficile* virulence.

## Main

Toxigenic bacteria possess diverse secretion systems to facilitate toxin release^1^. Structural aspects of these systems relate to the differing membrane and cell wall structures between Gram positive and Gram negative species, which need to be traversed in facilitating the release of cytoplasmically synthesized toxins. Gram-positive bacteria, which lack the Gram negative outer membrane, leverage two main pathways for protein secretion, namely the general secretory (Sec) pathway and the twin-arginine translocation (Tat) pathway^2^. Proteins using these systems contain an N-terminal secretion signal recognition sequence for membrane passage and release^3^.

In contrast to other bacterial toxins, large clostridial toxins (LCTs) lack identifiable secretion signals. LCTs form a family of bacterial exotoxins that exceed 200 kDa in size, and that inactivate host small GTPases to disrupt the actin cytoskeleton and promote host cell death^4,5^. In addition to *C. difficile* toxins TcdA and TcdB, members of the LCT family include *Clostridium novyi* toxin TcnA, *Clostridium perfringens* toxin TpeL and *Paeniclostridium sordellii* toxins TcsL and TcsH. Recent studies in *C. difficile*, *P. sordellii* and *C. perfringens*, suggest that LCT toxin secretion requires a conserved non-lytic holin-mediated transport mechanism^6–9^.

*C. difficile’s* Pathogenicity Locus (PaLoc) encodes the *tcdA* and *tcdB* genes and three accessory genes: *tcdR, tcdE* and *tcdC* (Fig. 1a)^10^. TcdR is an alternative sigma factor required for PaLoc gene expression, while TcdC has been proposed to negatively regulate transcription as an anti-sigma factor^11–15^. TcdE shares homology with bacteriophage holins and has been shown to promote toxin release^8,16^. Holins are small, bacteriophage-encoded membrane proteins that promote bacteriophage release. They oligomerize to form pores in the cytoplasmic membrane, supporting the export of bacteriophage-encoded endolysins which digest the peptidoglycan cell envelope and lead to cell lysis^17^.

**Fig. 1.**
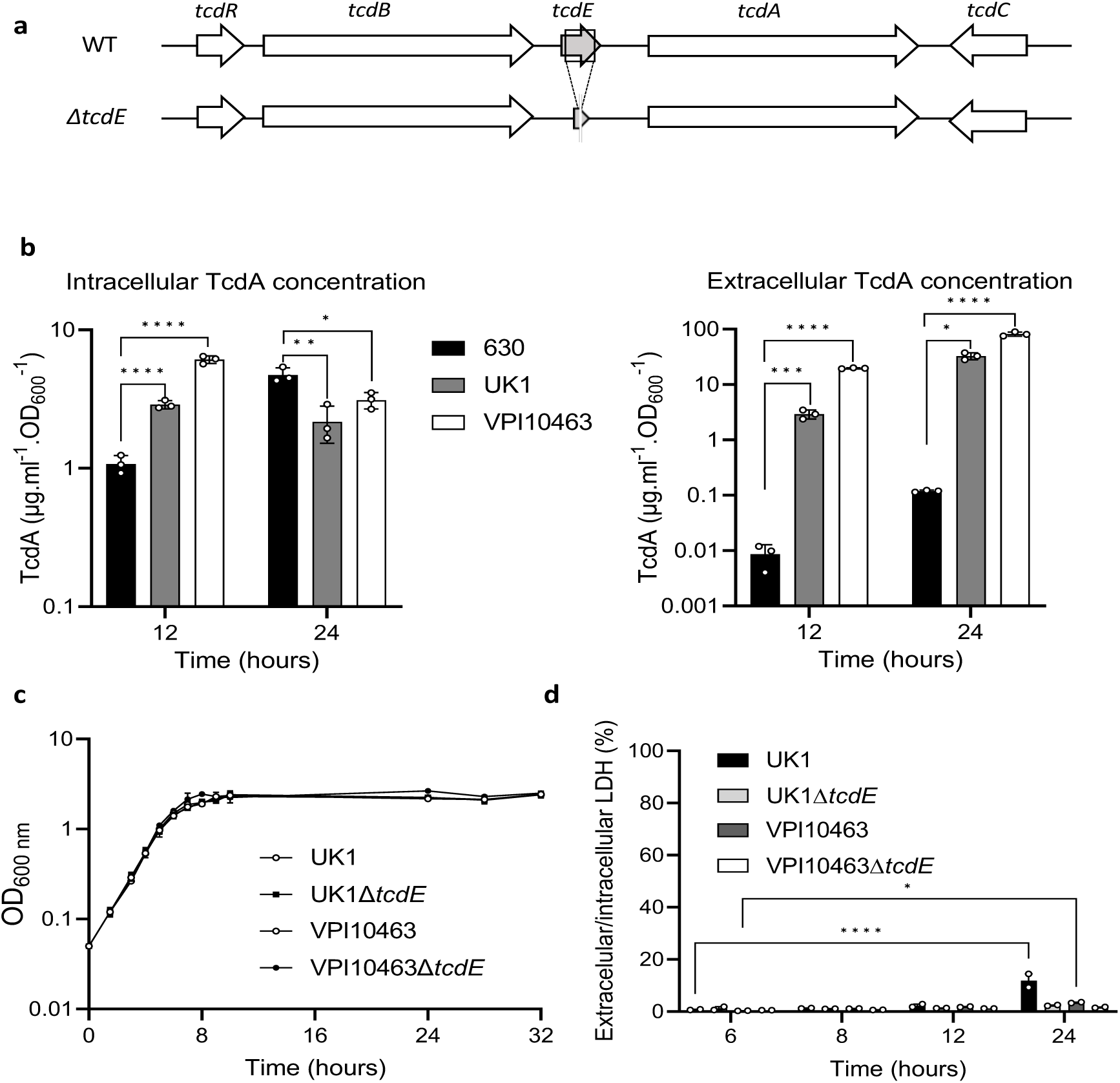
Toxin release independently of cell lysis in *C. difficile* VPI10463 and UK1. **a**. Schematic representation of the *tcdE* genetic environment and *tcdE* deletion (Δ*tcdE*). **b**. TcdA titers in extracellular and intracellular fractions of 630Δ*erm*, VPI10463 and UK1 strains after 12 and 24 hours of growth. The strains were grown in TY medium and TcdA was quantified using TcdA-ELISA. Means and SD are shown; n=3 independent experiments. \**P* ≤ 0.05, \*\**P* ≤ 0.01, \*\*\**P* ≤ 0.001 and *** *P* ≤ 0.0001 by a one-way ANOVA. **c**. Growth curves of VPI10463 and UK1 strains, and their respective Δ*tcdE* mutants in TY medium. Means and SD are shown; n=3 independent experiments. **d**. Ratio of the lactate dehydrogenase (LDH) activity in the supernatants and the cell lysates of VPI10463 and UK1 strains and their respective Δ*tcdE* mutants used as an indicator of autolysis. LDH activity was measured using the CytoTox 96 Non-Radioactive Cytotoxicity Assay (Promega). Means and SD are shown; n=2 independent experiments. \**P* ≤ 0.05 and *** *P* ≤ 0.0001 by a two-way ANOVA followed by a Dunnett’s multiple comparison test.

In *C. difficile,* non-lytic TcdE-dependent translocation of the PaLoc toxins^8^, in addition to lytic release in stationary phase via activity of the Cwp19 lytic peptidoglycan transglycosylase^18^, suggested that lytic and non-lytic mechanisms may coexist, and may function during different phases of cell growth and relative to stresses encountered *in vivo*^19^. Both systems occur in other toxigenic *Clostridia* that use holin-like proteins for LCT release, such as TpeL in *C. perfringens* and TcsL in *P. sordellii*^6–9^, and that also harbor Cwp19 homologs, further supporting functions of these systems in *Clostridial* toxin release. Within this framework, we hypothesized that the TcdE holin may mediate efficient release during periods of high-toxin expression, including in epidemic strains prone to producing high levels, with lytic mechanisms providing a default pathway integrated with physiologic and stress responses that induce pathogen cell lysis^16^.

We evaluated TcdE’s functions on toxin release *in-vitro* and *in-vivo* in the high-toxin producing and clinical strains VPI10463 and ribotype 027 epidemic strain UK1^20,21^. Both strains cause symptomatic infections in conventional and defined-association mouse models, resulting in severe disease and death^22,23^. We first compared intracellular and extracellular concentrations of TcdA in these strains and in the laboratory strain 630 at 12 and 24h of growth in tryptone yeast extract (TY) broth, a medium that supports high toxin expression^24, 25^. As expected, VPI10463 and UK1 produced higher toxins level of toxin than the 630 strain (Fig. 1b). Intracellular concentrations of TcdA were slightly lower at 12h in strain 630 but attained a comparable concentration with VPI10463 and UK1 by 24h. In contrast, extracellular concentrations of TcdA were 330 and 2,200-fold higher in UK1 and VPI10463, respectively at 12h, when compared to strain 630 (Fig. 1b). These findings illustrate that the highly virulent strains VPI10463 and UK1 release the majority of toxins produced, while toxin remained in the intracellular compartment of the less virulent and low toxin-producing 630 strain (Extended Data Fig. 1a). Comparable phenotypes in additional clinical strains were identified in the known high producer CD21-013 and low producer E25 (Extended Data Table 1; Extended Data Figs. 1B-C). These finding suggest that toxins first accumulate in the cytosol and are then secreted only upon reaching a certain threshold.

We generated in-frame *tcdE* deletion mutants to further evaluate TcdE’s functions in toxin secretion in VPI10463 and UK1 (Fig.1a, Extended data Fig.2). The wild-type and isogenic mutant strains grew similarly in TY media (Fig. 1c). Extracellular and intracellular lactate dehydrogenase (LDH) activity, a cytoplasmic enzyme used to monitor cell lysis, confirmed nominal cellular lysis between the wild-type and *tcdE* mutant strains in the first 12 hours of growth (Fig. 1d). However, the extracellular/intracellular ratio of LDH increased significantly at 24h in the parent UK1 strain, and to a lesser extent for VPI10463, but not in the *tcdE* mutants suggesting that TcdE activity contributed to cell lysis during late stationary phase.

**Fig. 2.**
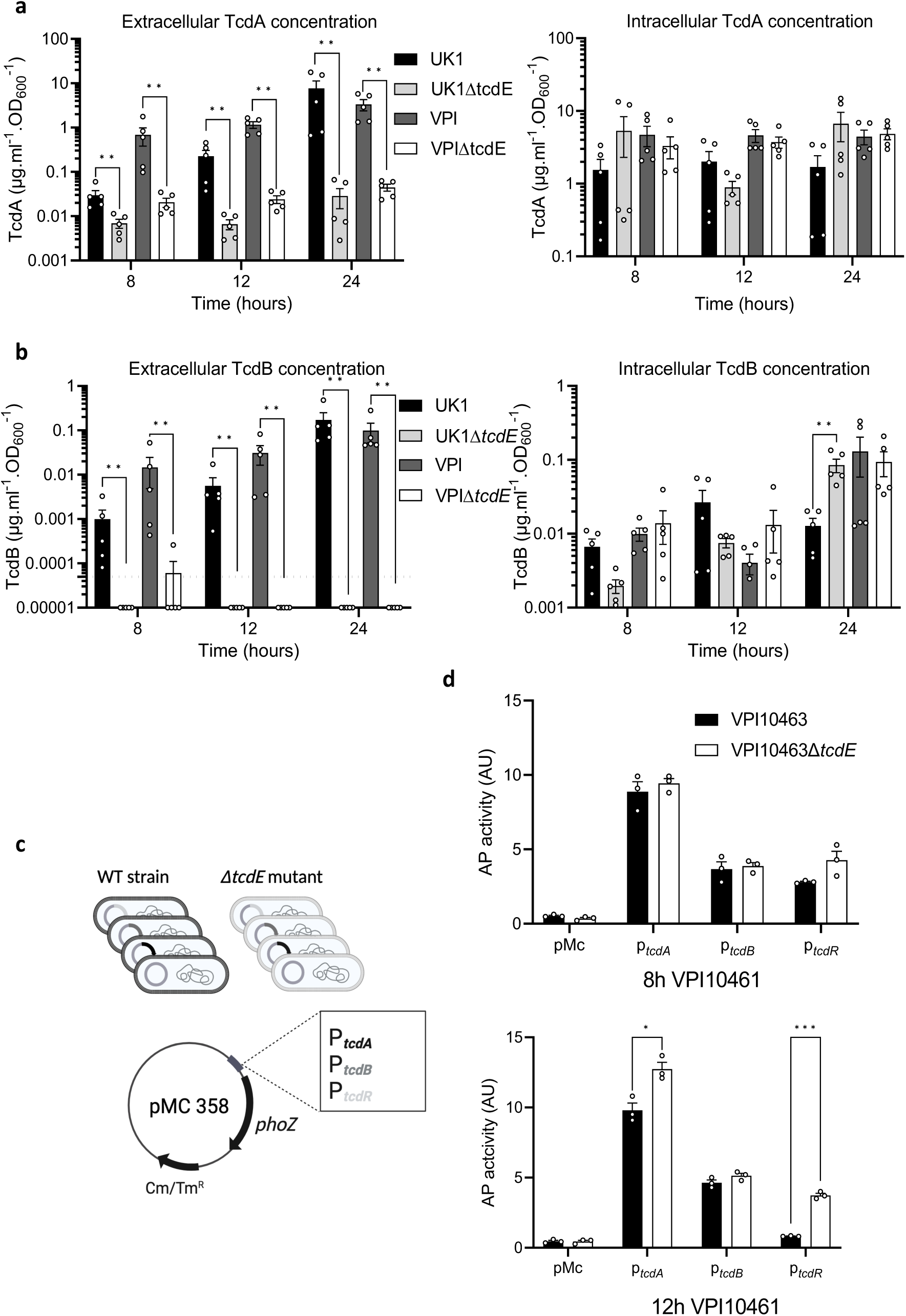
TcdE mediates TcdA and TcdB release in VPI10463 and UK1 strains *in vitro*. **a.** and **b.** TcdA (a) and TcdB (b) titers in extracellular (left panel) and intracellular (right panel) fractions of VPI10463 and UK1strains and their respective Δ*tcdE* mutants, after 8, 12 and 24 hours of growth. Strains were grown in TY medium, and toxins were quantified using TcdA-and TcdB-ELISA. Means and SEM are shown; n=5 independent experiments. ** p<0,01 by a Mann-Whitney test. Horizontal dotted line shows thresholds of detection. **c**. Schematic representation of transcriptional fusions constructions. Transcriptional fusions of promoter regions of approximately 500 bp of *tcdA*, *tcdB* or *tcdR* genes fused to the reporter gene *phoZ*, were introduced by conjugation into the VPI10463 wild type strain and the isogenic *ΔtcdE* mutants. **d**. Alkaline phosphatase (AP) activity of P*tcdA*::*phoZ*, P*tcdB*::*phoZ* and P*tcdR*::*phoZ* fusions expressed from pMC358 in VPI10463 and VPI10463Δ*tcdE*. Strains were grown in TY medium and samples assayed for AP activity were collected at 8 and 12 hours of growth. Means and SEM are shown; n=3 independent experiments. \**P* ≤ 0.05 and \*\*\**P* ≤ 0.001 by an unpaired t test.

Extracellular and intracellular toxin levels measured at 8, 12 and 24 h of growth showed dramatic reductions in extracellular TcdA and TcdB in the *tcdE* mutants of UK1 and VPI10463 (Fig. 2a and 2b). Strikingly, extracellular TcdB levels fell below the limit of detection in the *tcdE* mutants. Conversely, intracellular TcdA and TcdB accumulation remained comparable between the *tcdE* and the wild-type strains (Fig. 2a and 2b). To evaluate if *tcdE* deletion affected toxin gene transcription, plasmids carrying transcriptional fusions of the *tcdA, tcdB* and *tcdR* promoter regions with the alkaline phosphatase (AP) gene *phoZ*^26^ were transferred into the *tcdE* mutant and wild-type strains (Fig.2c). AP activities at 8h and 12h of growth were comparable in the VPI10463Δ*tcdE* and the isogenic wild-type strain, with a slight increase in *tcdR* and *tcdA* transcription at 12 h (Fig. 2c). The same findings occured with AP levels in the wild-type and isogenic Δ*tcdE* strains of UK1 carrying the transcriptional fusions (Extended data Fig. 4). qRT-PCR analyses of *tcdA*, *tcdB* and *tcdR* transcription in VPI10463 further showed no significant changes in PaLoc gene expression (Extended data Fig.3), confirming that *tcdE* deletion had no effects on toxin gene expression. These data highlight TcdE’s crucial role in toxin secretion in clinical epidemic strains, and to facilitate toxin release in a mechanism independent of cell-lysis (Fig. 1d).

**Fig. 3.**
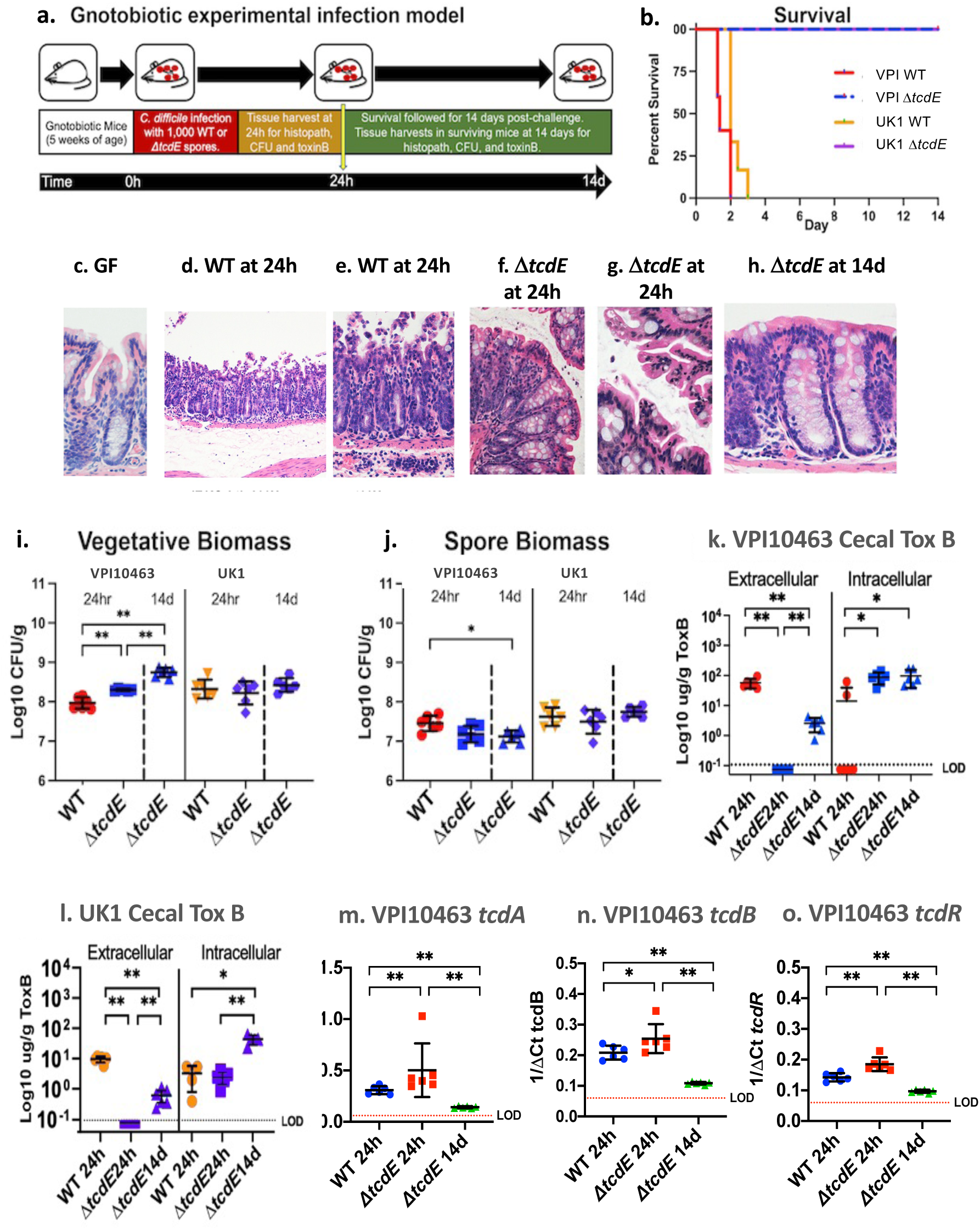
TcdE deletion rescues gnotobiotic mice from lethal *C. difficile* infection.: **a.** Experimental overview of the gnotobiotic infection model. **b.** Survival of mice challenged with wild-type VPI10463 (red), VPI10463*ΔtcdE* (blue), UK1 (orange) or UK1*ΔtcdE* (purple), n=8 mice per strain. **c-h.** Hematoxylin and eosin (H&E) sections of the colonic mucosa. **c.** Germ-free GF mouse at 400X magnification showing intact epithelium and normal lamina propria cellularity. **d.** Representative GF mouse infected with wild-type VPI10463 at 24h post-challenge, 100X magnification showing severe mucosal damage with transmural inflammation, tissue edema (asterix) and formation of pseudomembranes above the surface epithelium. **e.** Inset from panel d. at 200X magnification showing transmural neutrophilic infiltrates as well as immune cells in sub-mucosal blood vessels, at the bottom. **f.** Colon at 200X magnification of *ΔtcdE*-infected mouse showing limited surface epithelial ruffling but intact epithelium and with nominal inflammatory infiltrates at 24 hours post-challenge, in contrast to mice infected with the wild-type strain (panels d-e). **g.** 400X magnification of surface colonic epithelium from *ΔtcdE*-infected mouse showing focal ruffling of surface colonocytes but without formation of pseudomembranes. **h.** 400X magnification of colonic mucosa at 14 days in a surviving *ΔtcdE*-infected mouse showing intact colonic epithelium and limited lymphocytic infiltrates in the lamina propria. **i.** *C. difficile* vegetative biomass in cecal contents from mice infected with VPI10463 and UK1 or the isogenic *ΔtcdE*-mutant strains at 24h and from the Δ*tcdE*-infected mice at 14 days. Significance values for non-parametric Kruskal-Wallis test shown are **p=0.0022. **j.** *C. difficile* spore biomass in cecal contents from mice infected with VPI10463 and UK1 or the isogenic *ΔtcdE*-mutant strains at 24h and from the Δ*tcdE*-infected mice at 14 days. Significance values for non-parametric Kruskal-Wallis test shown are *p=0.012. **k.** Cecal extracellular and intracellular toxin B levels from mice infected with wild-type VPI10463 or the isogenic *ΔtcdE*-mutant strains at 24h and from the Δ*tcdE*-infected mice at 14 days. Dotted line with LOD indicates limit of detection. Significance values for non-parametric Kruskal-Wallis test shown are **p=0.0022 and *p=0.012. **l.** data comparable to panel k. for strain UK1. **m-o.** q rtPCR of cecal *tcdA, tcdB* and *tcdR* expression in mice infected with the wild-type or *ΔtcdE* VPI10463 strains at 24 hours and surviving *ΔtcdE* strain-infected mice at 14 days. Bars show mean and standard deviation. Kruskal-Wallis significance values as in panels k and l.

The ribotype 027 strain UK1 produces an additional toxin, binary cytolethal distending toxin (CDT), encoded by unlinked genes in the Cdt toxin locus, or CdtLoc ^27^. SignalP^28^ analyses identified putative Sec signal peptides in the N-terminus of the CDT subunits CdtA and CdtB, suggesting release by a holin-independent manner (Extended data Fig. 5a). We confirmed this finding in the UK1 *tcdE* strain which showed no impact on CDT secretion (Extended data Fig. 5b), indicating specificity of TcdE-dependent release for the TcdA and TcdB toxins.

To evaluate the impact of *tcdE* deletion on *C. difficile’s* virulence *in vivo*, 5-week-old germ-free mice were infected with the Δ*tcdE* mutants or the wild type strains (Fig. 3a). Mice infected with 1000 spores of wild-type VPI10463 rapidly succumbed to infection over 48-72 hours (Fig. 3b) with symptoms of lethargy, weight loss (extended data Fig. 6a), and diarrhea developing at 20 hours post-challenge. By 24 hours, infected mice demonstrated transmural inflammation with pseudomembrane formation and substantive mucosal edema when compared to the colon of germ-free mice (compare Fig 3c, with 3d-e). In contrast, all mice infected with the isogenic Δ*tcdE* strain survived infection (Fig 3b). Δ*tcdE*-infected mice demonstrated mild to sub-clinical symptoms of infection with no loss of weight (extended Fig. 6a). The colonic mucosa of Δ*tcdE-* infected mice demonstrated focal areas of mild apical epithelial ruffling at 24 hours post-challenge but without broad epithelial destruction and associated transmural inflammation (Fig 3f-g). By 14 days post-challenge, *ΔtcdE-*infected mice demonstrated intact colonic epithelium with mild lymphocytic infiltrates in the mucosa (Fig.3h). Mice infected with the wild-type and Δ*tcdE* mutant strains of UK1 similarly showed 100% rescue from lethal infection with the Δ*tcdE* mutant strain (Fig 3b). Clinically, mice infected with the Δ*tcdE* mutant of UK1 showed no weight loss (extended data Fig. 6b) and demonstrated comparable histopathologic findings in the colonic mucosa to mice infected with Δ*tcdE* mutant of VPI10463 (data not shown).

The vegetative biomass of VPI10463 *ΔtcdE* was 2-fold higher than that of the wild-type VPI10463 at 24 hours post-challenge and increased 5-fold from 24 hours to 14 days (Fig. 3i). In contrast, spore biomass remained equivalent between the strains, including at 14 days in the *tcdE* mutant (Fig.3j). In UK1-infected mice, vegetative and spore biomass remained comparable between the wild-type and the *ΔtcdE* strain over time (Fig. 3i and 3j). These data indicate that the *tcdE* strains have comparable or slightly better fitness to colonize the gnotobiotic mouse intestine than the corresponding wild-type strains.

At 24h of infection, while VPI10463-infected mice demonstrated high levels of TcdB toxin, levels fell below detectable thresholds in mice infected with the isogenic *tcdE* mutant (Fig. 3k). By 14 days post-infection, low toxin B levels were detectable in surviving *tcdE-*infected mice, suggesting release by a TcdE-independent mechanisms such as via cellular lysis, but remained 20-fold lower than the acute levels seen at 24h in mice infected with the wild-type strain (Fig. 3k). In contrast, intracellular TcdB levels were slightly increased in VPI10463 Δ*tcdE* at 24h post-infection when compared to the isogenic wild-type and remained constant over time in the mutant strain (Fig. 3k). Intra-and extracellular levels of TcdB in mice infected with the UK1 and UK1 Δ*tcdE* strains were comparable (Fig. 3l). To rule out effects of *tcdE* deletion *in vivo* on toxin gene expression, qRT-PCR of *tcdA*, *tcdB* and *tcdR* transcripts in VPI10463 (Fig 3m-o, respectively) showed elevated expression in the *ΔtcdE* mutant at 24h of infection, as compared to the wild-type strain. Expression levels in the *ΔtcdE* mutant fell 2 to 5-fold by 14 days in surviving *ΔtcdE* mutant-infected mice (Fig. 3o).

Our findings demonstrate the profound effects of *C. difficile’s* TcdE on host outcomes from infection, per its dominant role in facilitating extracellular toxin release. In contrast, release from alternate mechanisms, including cellular lysis via Cwp19 or other lytic mechanisms, was nominal per undetectable levels of extracellular toxin acutely in *tcdE* mutant-infected mice, and limited mucosal damage. By 14 days post-challenge, while extracellular toxin was detectable, it remained at levels 20-fold lower than seen at 24h in infection with the wild-type strains, suggesting that alternate mechanisms of toxin release play a limited role *in vivo,* for high toxin-producing strains^18^.

While the TcdE holin’s role in toxin release is clear, how it facilitates release remains an interesting question. In other systems, such as type 10 secretion systems (T10SS), holins have been shown to promote protein secretion through novel mechanisms that use a holin/endolysin pair^29^. In Gram negative species, the T10SS holins oligomerize to form pores in the cytoplasmic membrane, enabling the transport and release of a cytosolic endolysin into the periplasm. The peptidoglycan hydrolase activity of the endolysin locally permeabilizes the cell wall to allow specific protein substrates to be transported through the peptidoglycan layer and secreted^29^. The *Serratia marcescens* chitinases leverages this mechanism, via translocation of the endopeptidase endolysin ChiX into the periplasmic space via the holin ChiW^30^. Similarly, the secretion of the unusual typhoid AB-toxin of *Salmonella enterica typhi* requires peptidoglycan remodeling by muramidase activity of endolysin TtsA, which is believed to be translocated across the internal membrane by an as yet unidentified holin^31^. Similar to the *Clostridial* LCT loci, the Gram negative T10SS loci encode their secreted targets within the same genetic locus as the holin/endolysin pair^32^, suggesting potential for the TcdE holin to act in a comparable manner. Within the *C. difficile* PaLoc, most strains produce an endolysin remnant, designated TcdL^33^. However, an intact version of the endolysin has been identified in cryptic clades of *C. difficile* that carry a “Mono-Toxin B Paloc” ^34,35^. These findings raise the interesting question if *C. difficile* may use the TcdL remnant endolysin, or other endolysins located elsewhere in the genome, with TcdE to facilitate PaLoc toxin release.

Though LCTs use a holin/endolysin pair to traverse the peptidoglycan layer in *Clostridia*, how they cross the cell membrane remains ill-defined, given their lack of an N-terminal signal sequence. Recent studies by Saadat and Melville suggest that *C. perfringens’* LCT toxin TpeL can be directly transported by its TpeE holin using a charge zipper mechanism. After insertion into the membrane, TpeL induces folding and oligomerization of TpeE around it, to form a pore that facilitates subsequent secretion^7^. In the case of *C. difficile,* elucidation of the essential function of the TcdE holin in PaLoc toxin release opens opportunities to further resolve mechanisms of toxins transport through the cellular membrane and peptidoglycan layers, including requirements for local peptidoglycan remodeling by other PaLoc or chromosomal genes. Our findings also identify TcdE as a central target of vulnerability in *C. difficile’s* PaLoc toxin release, particularly during acute infections with high toxin-producing and epidemic strains, and support therapeutic interventions that reduce its expression and function *in vivo*.

## Methods

### Bacterial strains and cultures conditions

*C. difficile* and *Escherichia coli* strains used in this study are described in Extended DataTable 1. *C. difficile* strains were grown in a Freiter’s chamber (Jacomex) under anaerobic atmosphere (5% H2, 5% CO2, and 90% N2) in TY^24^ or Brain Heart Infusion (BHI, Difco) media. When appropriate, cefoxitin (Cfx, 25µg/ml) and D-cycloserine (Cs, 250 µg/ml) and thiamphenicol (Tm, 7.5 μg/ml) were added to the culture medium. *E. coli* strains were grown in LB broth, and when necessary, ampicillin (100 μg/ml) or chloramphenicol (15 μg/ml) was added to the culture medium. Growth curves were obtained by monitoring OD600 at each time point from an overnight culture diluted to a starting OD600 of 0.05. For quantitation of total biomass of *C. difficile*, mouse cecal contents were collected into pre-weighed Eppendorf tubes with 0.5mL of pre-reduced PBS with 40mM cysteine (Millipore-Sigma) as a reducing agent. Tubes were weighed after adding material and transferred into a Coy anaerobic chamber (Coy Labs) at 37°C for serial dilutions with plating to selective C. difficile CHROMID agar (Biomérieux) or Brucella agar (Becton Dickinson) for commensal quantitation. *C. difficile* colonies were counted at 48 hours of incubation. *C. difficile* spore preparations and counts were defined by exposing pre-weighed material to 50% ethanol for 60 minutes followed, by serial dilution and plating to *C. difficile* CHROMID agar. Vegetative cell biomass was calculated by subtracting the spore biomass from the total biomass and normalizing to the cecal mass.

### Plasmid and strain construction

All primers and plasmids used are listed in Extended Data Table 1 and 2. The *ΔtcdE* mutants in the *C. difficile* VPI10463 and UK1 backgrounds were obtained by allele-coupled exchange using the pMSR0 pseudo-suicide plasmid^36^. Briefly, homology arms of both up-and downstream locations of the target gene were amplified by PCR from genomic DNA of *C. difficile* VPI10463 and UK1 strains and purified PCR products were directly cloned into the pMSR0 vector using NEBuilder HiFi DNA Assembly (New England Biolabs). The pMSR0-derived plasmids containing the allele exchange cassettes of the target genes were transformed into *E. coli* strain NEB10β (New England Biolabs) and verified by sequencing. Plasmids were then transformed into *E. coli* HB101 (RP4) and transferred by conjugation into appropriate *C. difficile* strains. *C. difficile* transconjugants were selected on BHI agar supplemented with cefoxitin, D-cycloserine and thiamphenicol. Single cross-over integrants were selected based on the size of the colonies after restreak of the transconjugants on BHI with thiamphenicol. Colonies that underwent the second recombination event were then selected on BHI plates containing anhydrotetracycline (ATc: 100 ng/ml). Growing colonies were then tested by PCR for the presence of the expected deletion. For construction of promoter::phoZ constructs, promoter regions of *tcdA*, *tcdB* and *tcdR* of approximately 500 bp were amplified by PCR and cloned into the linearized pMC358 vector^26^.

### Alkaline phosphatase activity assays

*C. difficile* strains containing the *phoZ* reporter fusions were grown in TY medium at 37 °C in anaerobic conditions from an overnight culture diluted to a starting OD600 of 0.05 and harvested at 8 and 12hrs growth after inoculation. Cell pellets were washed with 0.5 ml of cold Wash buffer (10 mM Tris-HCl, pH 8.0, 10 mM MgSO4), centrifuged and resuspended in 750 µl Assay buffer (1 M Tris-HCl, pH 8.0, 0.1 mM ZnCl2). 500 µl of the cell suspensions were transferred into separate tubes and mixed with 350 µl of Assay buffer and 50 µl of 0.1% SDS, before vortexing for 1 min. Sample tubes were incubated at 37 °C for 5 min and then cooled on ice for at least 5 min. The assay starts by addition of 100 µl of 0.4% *p*NP (*p*-nitrophenyl phosphate in 1 M Tris-HCl, pH 8.0; Sigma-Aldrich) to each sample and incubation at 37 °C. A sample without cell was prepared as a negative control. Upon development of a light-yellow color, the alkaline phosphatase reaction was stopped by addition of 100 µl of stop solution (1 M KH2PO4) and placing the tubes on ice. Time elapsed (min) for the assay was recorded. Samples were then centrifuged at max speed for 5 min and absorbance for each sample was read at both OD420 and OD550. Units of activity were calculated and normalized to cell volume by using the following formula: (OD420 − (1.75 × OD550) × 1000)/(*t* (min) × OD600 × vol. cells (ml)).

### RNA extraction

Total RNAs were extracted from *C. difficile* strains grown in TY medium at 37 °C in anaerobic conditions up to 8, 12 and 24 hours. 10 ml of cultures were harvested and centrifuged (5000 rpm at 4°C) for 10 minutes. Supernatants were discarded and pellets immediately frozen and stored at −80°C. Pellets were then resuspended in 1mL of RNApro (MP biomedicals) and transferred into Matrix B tubes for lysis by FastPrep (MP biomedicals) with 3x 40‘’ shaking cycle at speed 6.5, separated by 2 min incubation in ice. Tubes were then centrifuged 10 min at 13000 rpm at 4°C (conditions used for the entire procedure). Half a volume of chloroform was thereafter added to the cell lysates and were vortexed for 5 sec. After 10 min centrifugation the upper phases were collected, and nucleic acids were precipitated by addition of 500µl of cold ethanol and stored at −20°C for at least 30 min. After a 15 min centrifugation, pellets were washed 3 times with 500µl of cold 75% ethanol following by 5 min centrifugation. Ethanol was carefully removed and pellets air dried at room temperature and resuspended in 60 µl of RNase free water. RNAs were then treated by DNase (Turbo DNA free Kit) according to the manufacturer’s recommendations.

### Reverse transcription and Real-time quantitative PCR

RNA (1µg) was incubated 10 min at 70°C with 1µg of hexamer oligonucleotide primers (p(DN)6, Roche) before adding 10 µl of 5X AMV RT buffer, 4 µl of dNTP (25mM), 1µl of RNasine (Promega) and 1µl of AMV RT enzyme (Promega) for a 50µl reaction volume. After 2h incubation at 37°C, the reaction was stopped by heat treatment (85°C) during 5 min. Real-time quantitative PCR was performed in 20μl reaction containing 20 ng of cDNAs, 10 μl of the SYBR PCR master mix (Life Technologies, Fisher) and 400 nM of *tcdA*, *tcdB*, or *tcdR* gene-specific primers (Extended Data Table 2). Amplification and detection were performed using a StepOne^TM^ instrument (Applied Biosystem). For each sample, the quantity of cDNAs of target genes was normalized to the quantity of cDNAs of the DNA polymerase III gene (*dnaF*). The relative change in gene expression was recorded as the ratio of normalized target concentrations (threshold cycle [ΔΔ*CT*] method)^37^

### Toxin ELISA

Overnight cultures of *C. difficile* strains were diluted in TY medium to a starting OD600 of 0.05 and grown at 37°C. 1 ml of culture was harvested after 8, 12 and 24 hours growth and centrifuged (5000 rpm at 4°C) for 5 min. Supernatants were collected and kept at −20°C. Cell pellets were washed in PBS 1X and kept at −20°C. For the Toxin A ELISA, PCG4.1 polyclonal antibodies (Bio-techne) were diluted at 4ng/ml in PBS and coated overnight on a Maxisorp plate to use as capture antibodies. The wells were then blocked with Superblock blocking buffer (Thermo Fisher Scientific). A range of purified toxin A (Merck) from 0 to 1 μg/ml used to perform a standard curve and dilutions of the samples were added to the wells. Detection antibodies anti-*C. difficile* toxin A coupled to HRP (LS-Bio) were then added at a 1:10000 dilution. The plate was developed by addition of TMB (Thermo Fisher Scientific) followed by a 30 min incubation and the reaction was stopped by addition of a 0.2M H2SO4 solution into the wells. The plate was read at a wavelength of 450nm with a Glomax plate reader (Promega). Toxin concentration in each sample was calculated using the standard curve and normalized by the optical density of the culture. The amount of toxin was normalized by the optical density of the culture. The same procedure is used for the Toxin B ELISA but N4A8 monoclonal antibodies (BBI solution) diluted at 4ng/ml in PBS was used for the capture antibodies and the T4G1 monoclonal antibodies previously coupled to biotin were used as detection antibodies (BBI solution, 1:10000 dilution) with streptavidin HRP (Thermo Fisher Scientific). For the CDT ELISA, chicken *C. difficile* binary toxin subunit B capture and detection antibodies (MyBiosource) were used following the supplier’s instructions.

### Lactate deshydrogenase (LDH) assays

1ml of culture aliquots were taken and pelleted at indicated time points by centrifugation for 5 min at 5000 rpm at 4°C. Supernatants were collected and cell pellets were resuspended in 1 ml of PBS. Resuspended pellets were transferred in Matrix B tubes for lysis by FastPrep (MP biomedicals) with 2×40’’shaking cycles at speed 6.5, separated by 2 min incubation in ice. LDH activity in the supernatants and the cell lysates was then assessed using the CytoTox 96 Non-Radioactive Cytotoxicity Assay (Promega) by following the manufacturer’s recommendations. Optical density was read at 490nm on a Glomax plate reader (Promega). The ratio of supernatant to total LDH activity normalized by the cell density of the samples was used as an indicator of autolysis.

### Germ-free mouse infection studies

Animal studies were conducted in negative pressure BL-2 gnotobiotic isolators (Class Biologically Clean, Madison, WI)^38^ under an approved institutional IACUC protocol. Equal ratios of 6 week-old male and female gnotobiotic mice were singly housed and challenged with 1×10^3^ of wild-type or Δ*tcdE C. difficile* spores. Progression of disease was assessed via body condition scoring^39^. Mice were sacrificed at a BCS of 2-, or at defined timepoints at either 24 hours or 14 days post-*C. difficile* challenge. Cecal contents were collected for functional studies including CFU enumeration, toxin ELISA, and qRT-PCR. The gastrointestinal tract and internal organs were fixed in zinc-buffered formalin (Z-FIX, Thermo-Fisher, Waltham, MA) for histopathologic assessment.

### Histopathological analyses

Formalin-fixed gut segments from germ-free or infected mice were paraffin embedded and 5mm sections cut for staining with hematoxylin and eosin (H&E; Thermo-Fisher, Waltham, MA) as described^40^. Slides were visualized under a Nikon Eclipse E600 microscope (Nikon, Melville, NY) to assess epithelial damage per cellular stranding and vacuolation, the nature of Inflammatory infiltrates, mucosal erosions, and tissue edema. Lumenal neutrophils were quantified by a Pathologist by evaluating ten 400X high powered fields (HPFs) across at least 3 colonic sections per mouse. Neutrophils were identified by presence of segmented nuclei and pale to finely granular cytoplasm.

### Statistical analysis

All statistical tests were performed in GraphPad Prism. A p value <0.05 was considered significant. For gnotobiotic mouse studies, survival studies were evaluated in Prism 9.0 (GraphPad, San Diego, CA) using the Mantel-Cox for assessment of significant differences between wild-type and Δ*tcdE* mutant strains. Vegetative and spore biomass, and toxin B levels from mice were analyzed in Prism 9.0 (GraphPad, San Diego, CA) for visualization. Significant differences among groups were evaluated by non-parametric Kruskal-Wallis ANOVA and Dunn’s post-test. A p value ≤0.05 was considered significant.

## Acknowledgments

This work was funded by the Institut Pasteur and the “Integrative Biology of Emerging Infectious Diseases” (LabEX IBEID) funded in the framework of the French Government’s “Programme Investissements d’Avenir” to B.D, the ANR-20-CE15-0003 (Difficross) to J.P, and grants R01AI153605, R03AI174158, and P30DK34854 from the National Institute of Health in the United States, and a capital grant from the Massachusetts Life Science Center (MLSC) to LB.

## Author information

These authors contributed equally: Nicholas V DiBenedetto and Marine Oberkampf

## Authors and affiliations

Massachusetts Host-Microbiome Center, Dept. Pathology, Brigham and Women’s Hospital, Harvard Medical School, Boston, MA, USA

Lynn Bry, Nicholas V DiBenedetto, Laura Cersosimo and V Yeliseyev Institut Pasteur, Université Paris-Cité, UMR-CNRS 6047, Laboratoire Pathogenèse des Bactéries Anaérobies, F-75015 Paris, France Marine Oberkampf and Bruno Dupuy Institute for Integrative Biology of the Cell (I2BC), CEA, CNRS, Université Paris-Saclay, Gif-sur-Yvette, France Johann Peltier

## Contributions

B.D., J.P. and L.B contributed to the experiment design and interpreted all of the results. M.O., J.P. and B.D. created bacterial strains and performed *in vitro* assays (AP, LDH and toxins) and QRT-PCR. N.V.D., LC and V.Y carried out mice experiments and all assays from cecal samples such as bacterial numeration, toxin assays and qRT-PCR. B.D. B.D., J.P. and L.B contributed to writing the first draft of the manuscript and all of the authors commented on manuscript drafts

## Corresponding Author

Correspondence to Bruno Dupuy

## Ethics declaration

All studies were conducted under an approved institutional IACUC protocol. Mice were singly housed for all studies and fed *ad libitum* autoclaved LabDiet 5021 Autoclavable Mouse Breeder Diet (PMI Nutrition International, St. Louis, MO).

## Additional information

None.

## Extended data

Extended data Table 1 and 2, Extended data Figures 1 to 6.

---

**Extended data Fig 1:**
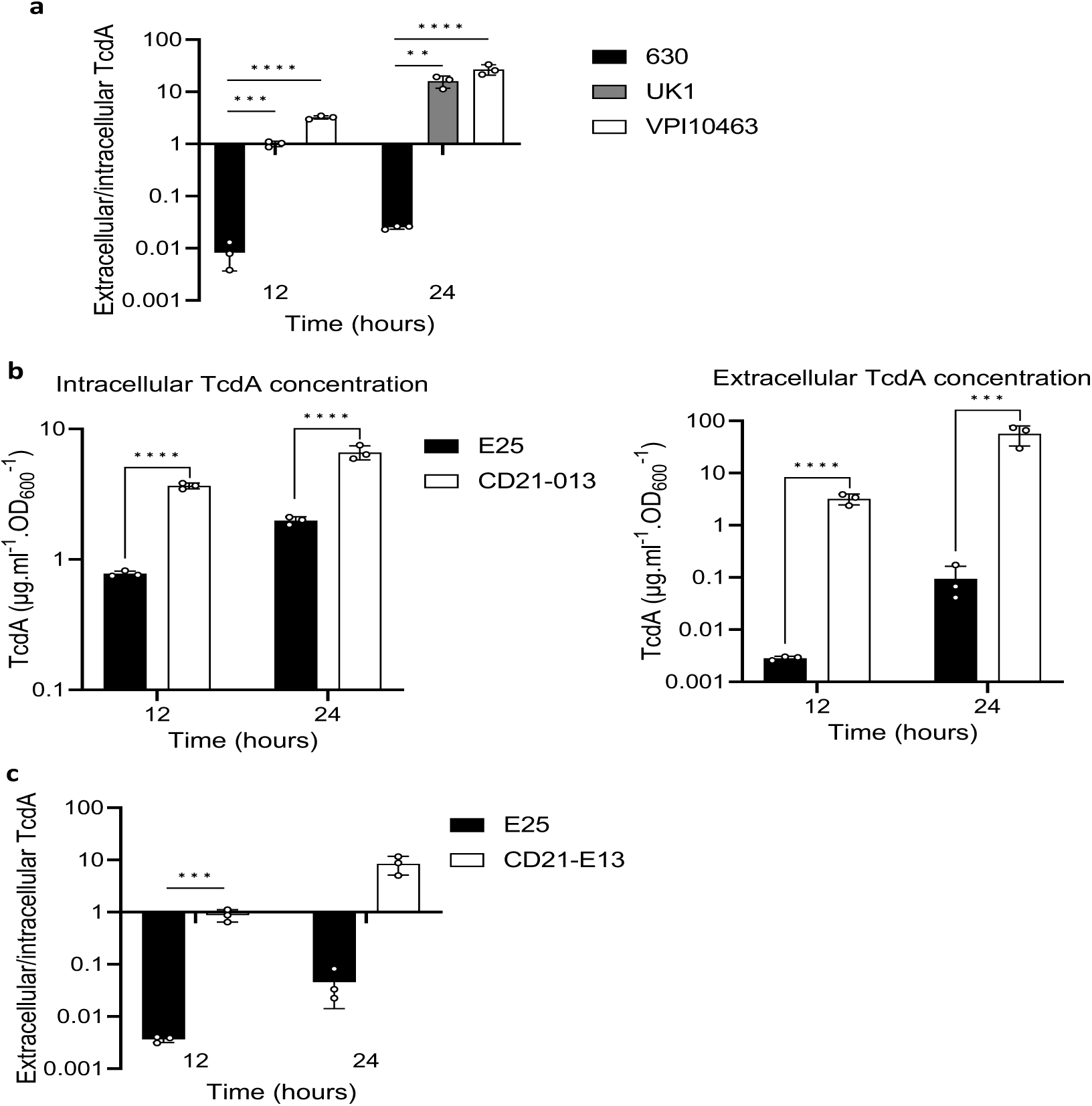
TcdA is actively secreted in high toxin producer strains. **a.** extracellular/intracellular ratio of TcdA from 630Δ*erm*, UK1 and VPI10463 strains. The strains were grown in TY medium and TcdA concentrations from the supernatant and the intracellular content were quantified using a TcdA-ELISA. Means and SD are shown; n=3 independent experiments. \*\**P* ≤ 0.01, \*\*\**P* ≤ 0.001 and *** *P* ≤ 0.0001 by a one-way ANOVA. **b.** TcdA titers in extracellular and intracellular fractions of E25 and CD21-013 strains after 12 and 24 hours of growth. The strains were grown in TY medium and TcdA was quantified using TcdA-ELISA. Means and SD are shown; n=3 independent experiments. \*\*\**P* ≤ 0.001 and *** *P* ≤ 0.0001 by a one-way ANOVA. **c.** extracellular/intracellular ratio of TcdA from E25 and CD21-013 strains. The strains were grown in TY medium and TcdA concentrations from the supernatant and the intracellular content were quantified using a TcdA-ELISA. Means and SD are shown; n=3 independent experiments. \*\*\**P* ≤ 0.001 by a one-way ANOVA.

**Extended data Fig.2:**
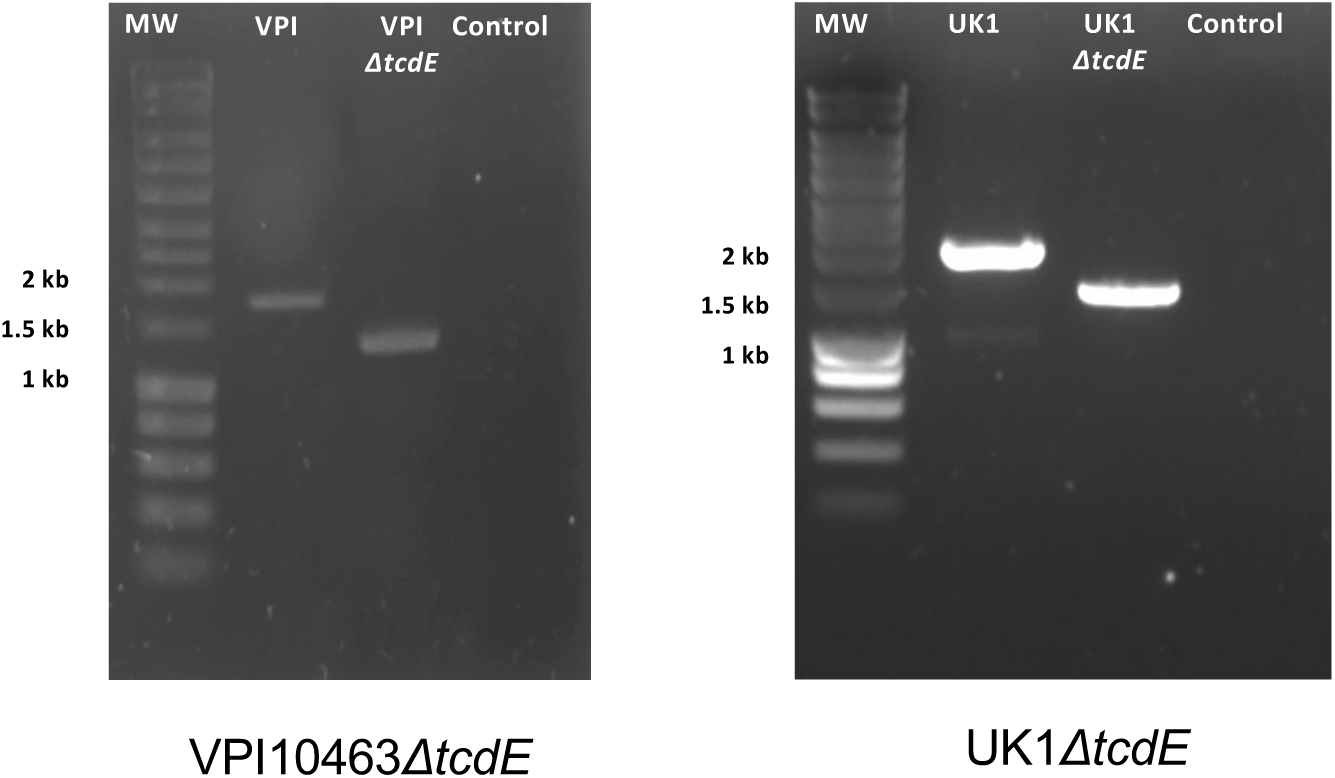
PCR verification of the *tcdE* deletion mutants in both *C. difficile* VPI10463 and UK1 strains.

**Extended data Fig 3:**
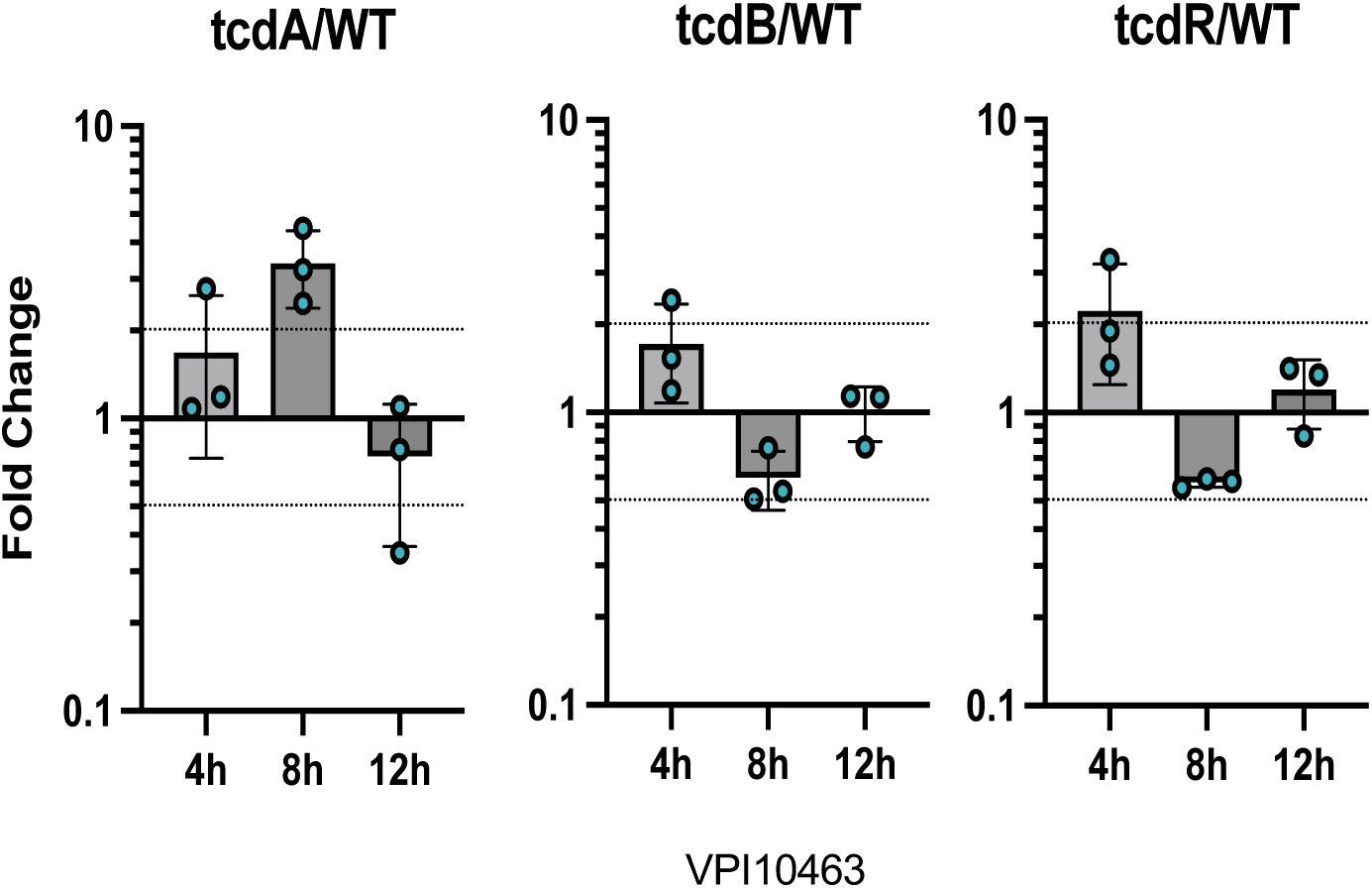
Transcript abundance of *tcdA*, *tcdB* and *tcdR* quantified by qRT-qPCR. *C. difficile* strains were grown in TY medium. Samples were collected after 4, 8 and 12 hours of growth. Means and SD are shown; n=3 independent experiments.

**Extended data Fig 4:**
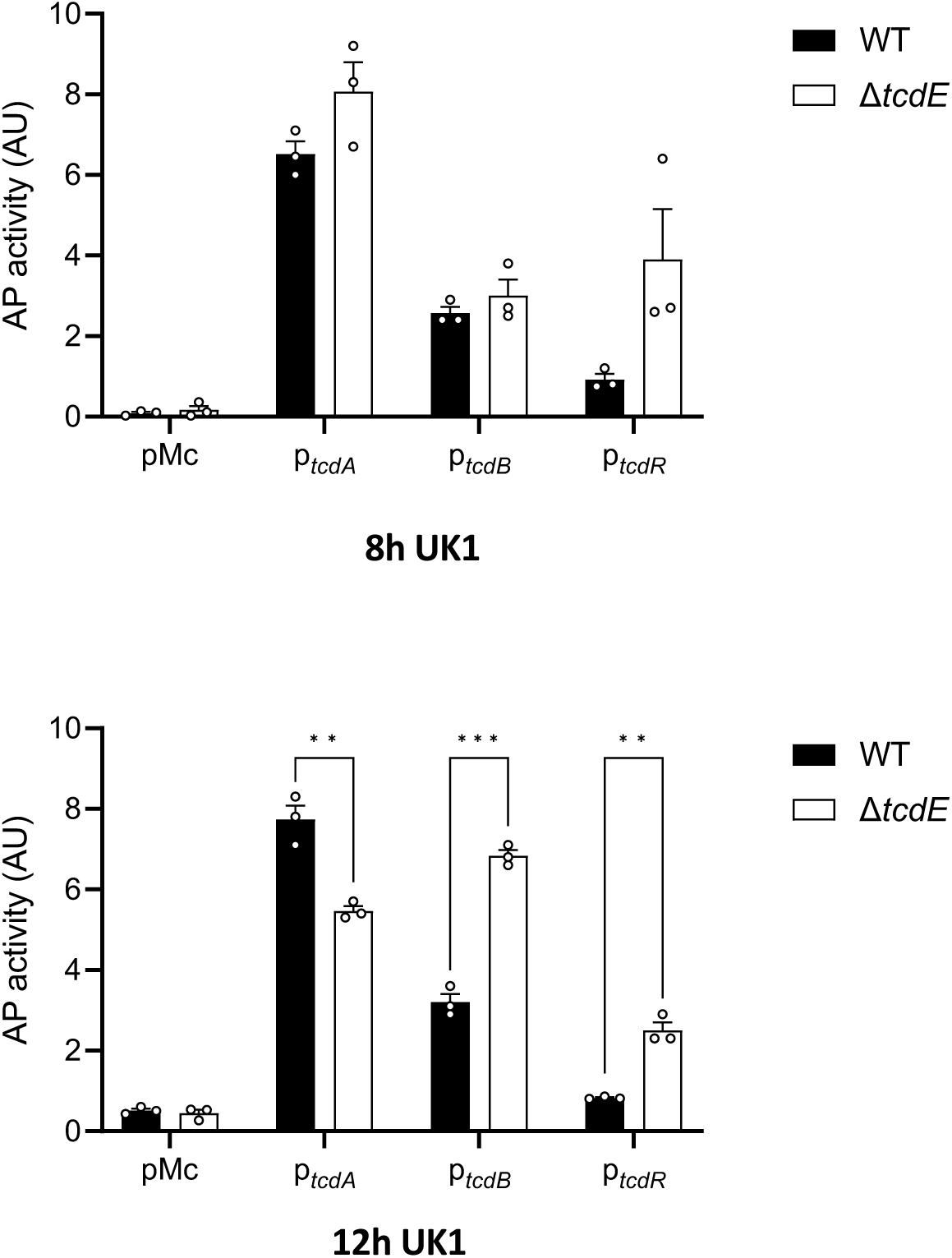
Alkaline phosphatase (AP) activity of P*tcdA*::*phoZ*, P*tcdB*::*phoZ* and P*tcdR*::*phoZ*, expressed from a plasmid in UK1 and UK1Δ*tcdE*. Strains were grown in TY medium and samples assayed for AP activity were collected at 8 and 12 hours of growth. Means and SEM are shown; n=3 independent experiments. \*\**P* ≤ 0.01 and \*\*\**P* ≤ 0.001 by an unpaired t test.

**Extended data Fig 5:**
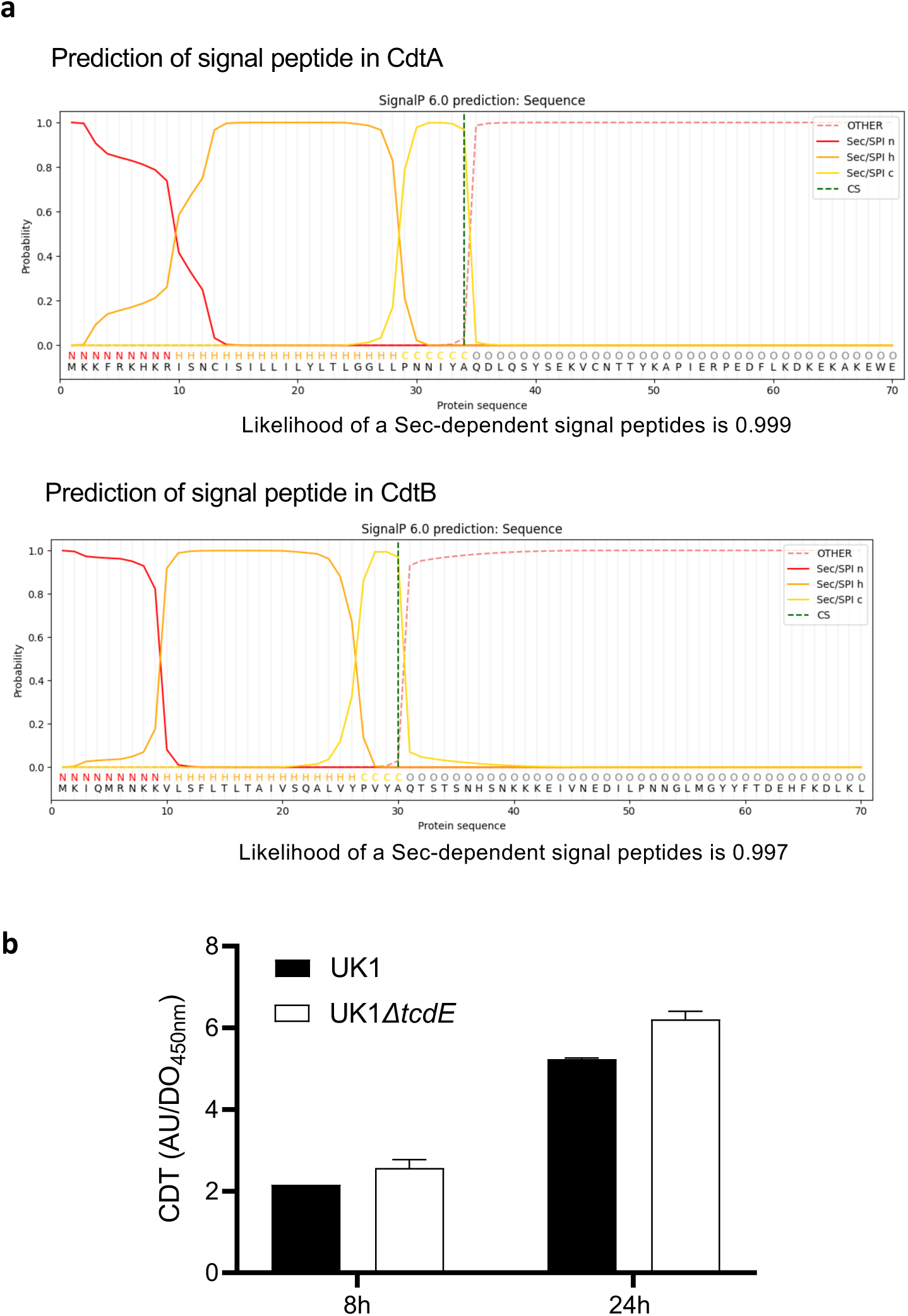
**a**. Prediction of a signal peptide from CdtA and CdtB amino acid sequences using SignalP 6.0^28^. **b**. Binary toxin (CDT) levels in extracellular fractions of the UK1 and UK1 Δ*tcdE* strains after 8 and 24h of growth. Strains were grown in TY medium, and CDT was quantified using CDT-ELISA. Means and SD are shown; n=3 independent experiments.

**Extended data Fig 6:**
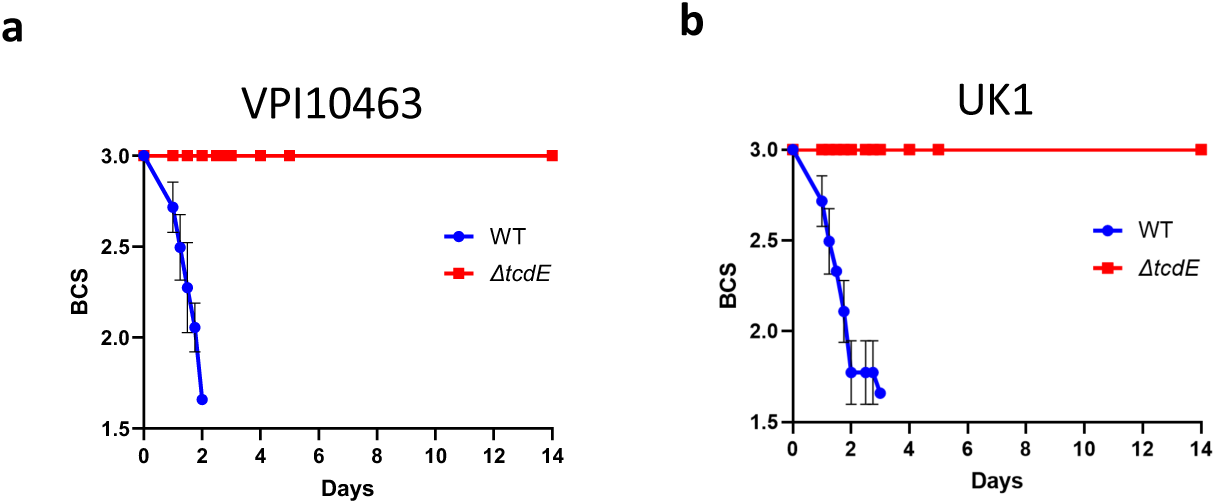
**a**. Body condition scores of mice infected with VPI10463 strain and its respective *tcdE* mutant. **b**. Body condition scores of mice infected with UK1 strain and its respective *tcdE* mutant.

**Extended data Table 1.** Strains and plasmids used in this study.

| Strain | Genotype | Origin |
| --- | --- | --- |
| *E. coli* |  |  |
| NEB-10 beta | $\Delta(ara-leu)$ 7697 *araD139 fhuA* $\Delta lacX74$ *galK16 galE15* *e14-* $\phi 80dlacZ\Delta M15$ *recA1 relA1 endA1 nupG rpsL* (Str<sup>R</sup>) *rph spoT1* $\Delta(mrr-hsdRMS-mcrBC)$ | New England Biolabs |
| HB101 (RP4) | *supE44 aa14 galK2 lacY1* $\Delta(gpt-proA)$ 62 *rpsL20* (Str<sup>R</sup>) *xyl-5 mtl-1 recA13* $\Delta(mcrC-mrr)$ *hsdS<sub>B</sub>* (r<sub>B</sub>-m<sub>B</sub>-) RP4 (Tra<sup>+</sup> IncP Ap<sup>R</sup> Km<sup>R</sup> Tc<sup>R</sup>) | Laboratory stock |
| EC1370 | HB101(RP4) carrying pDIA7080 | This work |
| EC1384 | HB101(RP4) carrying pDIA7090 | This work |
| *C. difficile* |  |  |
| 630$\Delta erm$ | 630 $\Delta ermB$ | Laboratory stock<sup>1</sup> |
| UK1 |  | Laboratory stock<sup>2</sup> |
| VPI10463 |  | Laboratory stock<sup>3</sup> |
| E25 |  | Laboratory stock<sup>4</sup> |
| CD21-013 |  | Laboratory stock<sup>5</sup> |
| CDIP1688 | VPI10463 $\Delta tcdE$ | This work |
| CDIP1855 | VPI10463 carrying pMC358 | This work |
| CDIP1920 | VPI10463 $\Delta tcdE$ carrying pMC358 | This work |
| CDIP1753 | UK1 $\Delta tcdE$ | This work |
| CDIP1838 | UK1 carrying pMC358 | This work |
| CDIP1854 | UK1 $\Delta tcdE$ carrying pMC358 | This work |
| CNRS_CD166 | VPI10463 carrying p112 | This work |
| CNRS_CD170 | VPI10463 $\Delta tcdE$ carrying p112 | This work |
| CNRS_CD167 | VPI10463 carrying p113 | This work |
| CNRS_CD171 | VPI10463 $\Delta tcdE$ carrying p113 | This work |
| CNRS_CD169 | VPI10463 carrying p115 | This work |
| CNRS_CD172 | VPI10463 $\Delta tcdE$ carrying p115 | This work |
| CDIP1834 | UK1 carrying p112 | This work |
| CDIP1850 | UK1 $\Delta tcdE$ carrying p112 | This work |
| CDIP1837 | UK1 carrying p113 | This work |
| CDIP1853 | UK1 $\Delta tcdE$ carrying p113 | This work |
| CDIP1836 | UK1 carrying p115 | This work |
| CDIP1852 | UK1 $\Delta tcdE$ carrying p115 | This work |
| **Plasmid** |  |  |
| pMSR0 | Allele exchange in *C. difficile* UK1 and VPI10463 | Peltier et al<sup>6</sup> |
| pMC358 | Promoterless *phoZ* | Edwards et al<sup>7</sup> |
| pDIA7080 | pMSR0 derivative for *tcdE* deletion in VPI10463 | This work |
| pDIA7090 | pMSR0 derivative for *tcdE* deletion in UK1 | This work |
| p112 | pMC358 derivative for P<sub>tcdR</sub>-*phoZ* fusion | This work |
| p113 | pMC358 derivative for P<sub>tcdB</sub>-*phoZ* fusion | This work |
| p115 | pMC358 derivative for P<sub>tcdA</sub>-*phoZ* fusion | This work |

1. Hussain, H.A., *et al.* (2005). *J Med Microbiol* 54, 137-41. 2. Killgore GTA, *et al.* (2008). *C. J Clin Microbiol* 46:431–437. 3. Hammond G.A and Johnson JL. (1995 ).*Micro. Patho.* **19**:4, 203-213 4. Kurka H *et al.*, (2014). *PloS One* 9:1, e86535 5. Anjou C *et al.*, *in preparation* 6. Peltier, J. *et al.* (2020) *Commun Biol* **3**, 718. 7. Edwards, A.N. *et al.* (2015). *Anaerobe* **32**, 98-104.

**Extended data Table 2.**
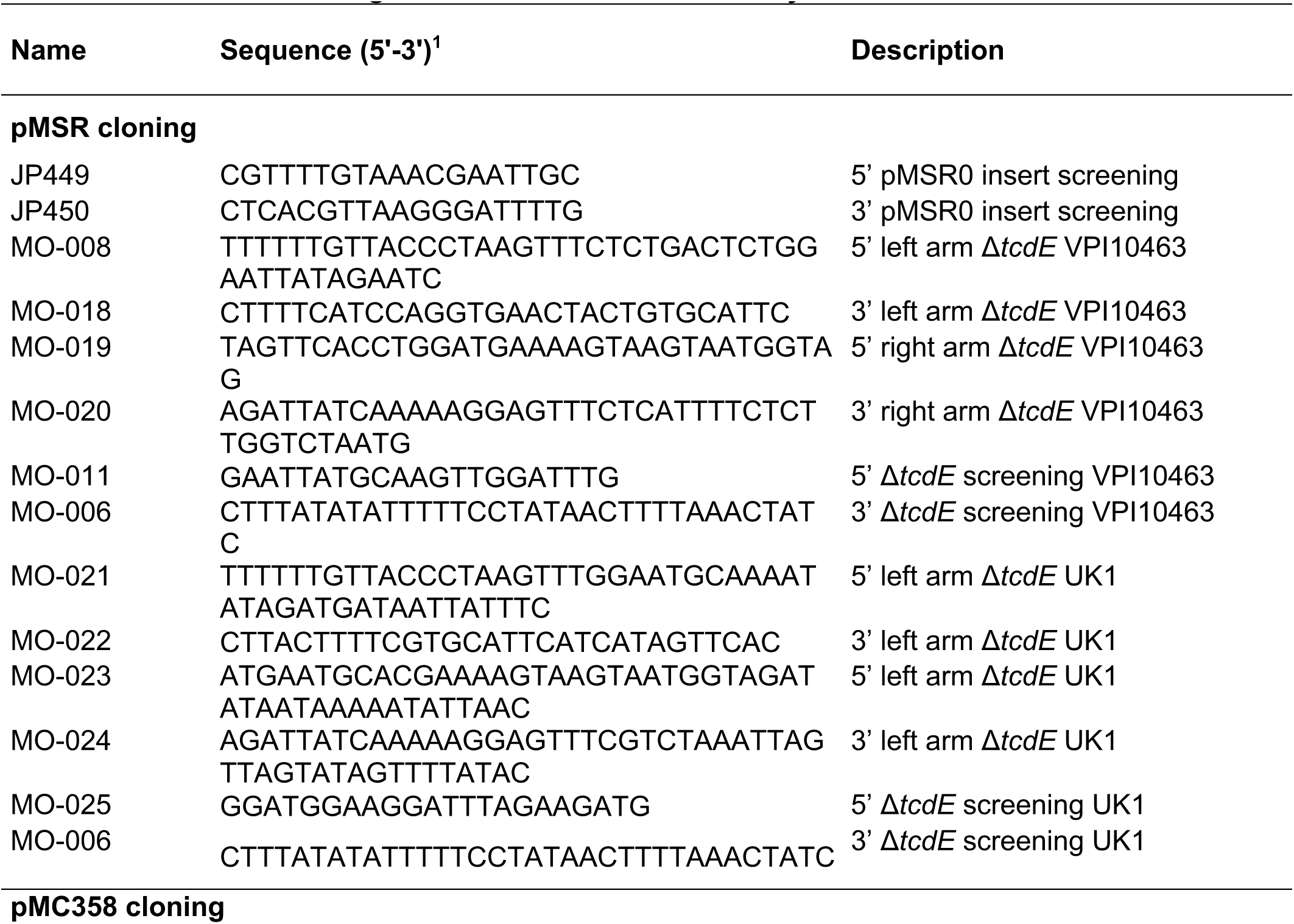
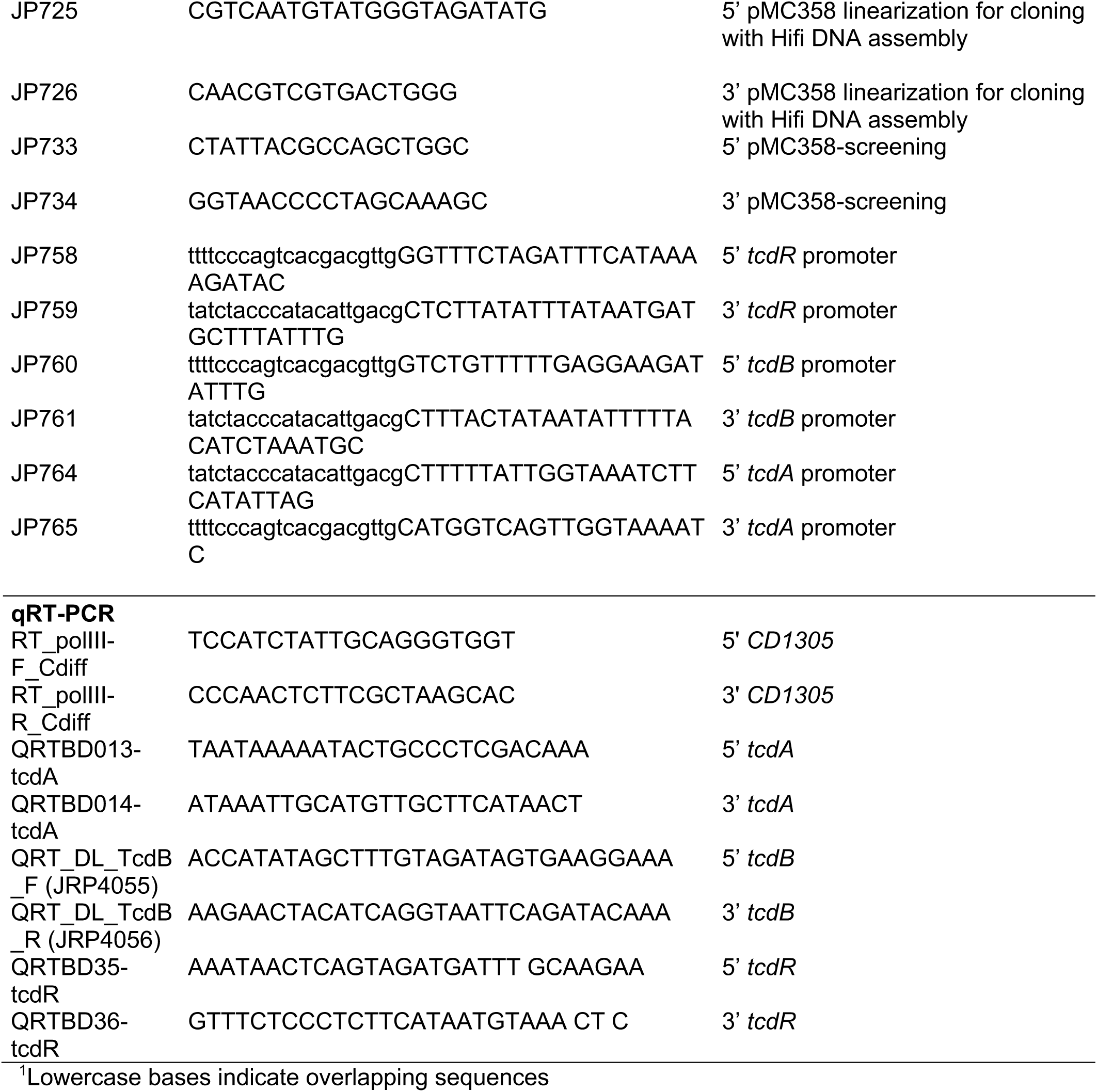
Oligonucleotides used in this study. Name.

